## Supporting Information for "Genetic underpinnings of chills from art and music"

**This PDF file includes:**
Supplementary Text
Supplementary Figure S1
Supplementary Figure S2
Supplementary Figure S3
Supplementary Table S1
Supplementary Table S2
Supplementary Table S3
Supplementary Table S4
Supplementary References

### Primary measures

Similar to previous studies(1,2), chills were selected as a trait of interest as individuals show chills while experiencing intense, pleasant emotional responses from diverse art forms. Self-reported chills correlate with objective chills(2–5), measured by physiological changes such as skin conductance, providing an easily assessable state with clear observable correlates(6). For example, across different experimental settings, the overlap between self-reported chills, as measured by button presses and skin conductance responses, varies between 46%(7) and 73%(3), with the highest correlations between chills measured by a combination of button presses and galvanic skin responses and self-reported chills of .90(8). In the present study, proneness to aesthetic chills was measured by the NEO questionnaire item: “wanneer ik een gedicht lees of naar een kunstwerk kijk, voel ik soms een koude rilling of een golf van opwinding” (English translation: “Sometimes when I am reading poetry or looking at a work of art, I feel a chill or wave of excitement”)(9). As in Bignardi et al.(10), this item was selected since it captures individual differences highly shared among cultures(9), it has been suggested to be a “universal emotional experience”(9), and to represent one of a few possible “vectors of biological variation common to all humans”(11) capturing meaningful inter-individual personality differences. Further, it is the only item assessing proneness to chills from art that has been shown to be heritable(10). We also note that this item can predict self-reported and objective chills from other art domains, such as music ($r$ = .29(8)) and correlates ($r$ ~ .10) with the strength of the functional connections between different cortical resting state networks(12). Proneness to music chills was assessed by the Barcelona Music Reward Questionnaire (BMRQ): “Soms krijg ik kippenvel als ik naar een liedje luister dat ik mooi vind” (English translation: “I sometimes feel chills when I hear a melody that I like”)(13). Although we are unaware of previous item-specific analysis conducted on this item, a composite score including the chill item has been shown to be heritable(14) and to correlate with self-reported and objectively assessed music chills(15), task-based functional cortico-subcortical connectivity(16), and white matter microstructure differences(17).

### Quality control of genotypes

Information about quality control of genotyped data and samples, and genetic imputation, can be found at https://wiki.lifelines.nl/doku.php?id=ugli. We performed additional SNP quality control in PLINK(18) v1.90b6.10on imputed data. SNPs were excluded if they had an imputation quality INFO score < 0.8 or a Minor Allele Frequency (MAF) < .01. Next, imputed data from release 1 (version 2) and 2 (version 2) were merged, keeping only SNPs available in both batches, and filtered for Hardy-Weinberg equilibrium (p <.0001).

### Descriptives

Within the sample of 35,114 people, proneness to aesthetic chills displayed a mean of 2.82 ($SD$ = 1.10) and a median of 3 and proneness to music chills a mean of 3.96 ($SD$ = 0.93) and a median of 4. For both items, scores ranged across the entire spectrum (i.e. 1 to 5; “strongly disagree” to “strongly agree”). Mean and median differences of aesthetic and music chills were significant; two-sided paired $t$ and Wilcox signed-ranked tests, t(35113) = -193.83, $p < 2.2\times10^{-16}$; V = 8824678, $p < 2.2\times10^{-16}$, respectively. This indicates that individuals tend to be more prone to music chills than aesthetic chills. Following reference(19), we then computed skewness as:

$$skew = \frac{\frac{1}{n}\sum_{i}^{N}(y_i - \bar{y})^3}{(\frac{1}{n}\sum_{i}^{N}(y_i - \bar{y})^3)^{\frac{3}{2}}} \tag{1}$$

where $y_i$ is the phenotypic score for individual $i$. The base R code to compute skewness can be found here: https://github.com/cran/moments/blob/master/R/skewness.R. Proneness to aesthetic chills displayed very little skewness (positively skewed, skew = 0.09), while proneness to music chills displayed substantial negative skewness (skew = -1.00, flat tail at the left side of the distribution).

Since proneness to music chills was substantially skewed, all further analyses reported in the Supplementary Text were carried out on fully adjusted two-stage ranked normalised data, when not otherwise specified (see below for details). Analyses in the main text were also carried out on transformed data. The main results for non-adjusted data are reported in Supplementary Table S1.

**Associations with demographics**

To assess potential effects of age and sex, we fit a linear model where proneness to chills (raw scores) was regressed on age and sex. To account for sample relatedness, the linear model was fit using the R package lavaan(20), which provides robust estimates of uncertainty around point estimates. Both proneness to aesthetic and to music chills were higher in women than men, with an average unit increase of $b$ = .43 (95% CI [.41; .45] ; $p$ < .001) and $b$ = .24 (95% CI [.22; .26]; $p$ < .001) from men to women ($b$ are unstandardised). In line with previous findings(10), age positively predicted proneness to aesthetic chills ($b$ = .009, 95% CI [.008; .010]; $p$ < .001) but negatively predicted proneness to music chills ($b$ = -.010, 95% CI [-.010; -.009]; $p$ < .001), suggesting different trajectories across different modalities over the lifespan. Since proneness to music chills was substantially skewed, we also repeated the analysis on the inversed-rank transformed and normalised data. Inversed ranked transformation followed:

$$y_i = \Phi^{-1}\left(\frac{R_i - .5}{N}\right) \tag{2}$$

where $\phi^{-1}$ is the inverse cumulative distribution function, and $R_i$ is the rank of the raw score $y$ value for individual $i$. Conclusions from this analysis were unchanged, with sex-effect being β =.19 (95% CI [.18; .20]; $p < 2.2\times10^{-16}$) and β = .12 (95% CI [.11; .13]; $p < 2.2\times10^{-16}$) and age-effect being β =.10 (95% CI [.09; .11]; $p < 2.2\times10^{-16}$) and β = -.13 (95% CI [-.14; .-11]; $p < 2.2\times10^{-16}$) for aesthetic chills and music chills, respectively (β are standardised).

**Fully-adjusted two-stage rank normalisation procedure**

Given that the Restricted Maximum Likelihood (REML) estimator can be sensitive to departures from normality, all results reported in the main text were derived from fully adjusted two-stage ranked normalised transformed data. Briefly, covariates (i.e., age, sex, genotyping array, and the first 10 Genomic principal components (PCs)) were regressed from the two phenotypes $Y_1$ and $Y_2$. Then, phenotypes were transformed following the inverse-rank transformation outlined in equation 2. It has been shown that this second step may introduce undesirable statistical properties, such as reintroducing effects of covariates(21). Therefore, following prior work(21,22), we regressed covariates once more from the transformed data. Final residuals were scaled. We note that residualisation and transformation were done with the reduced sample of 15,606 genotyped participants included for analysis. For comparison, GREML-based results obtained from raw scores, including covariate in the models, can be found in Supplementary Table S1. Code to apply the fully-adjusted two-stage rank normalisation procedure was adapted from de Hoyos et al.(22) and can be found following the link below:
https://github.com/ldehoyos/ASD_heterogeneity_grmsem_2024/blob/main/01_phenotypes/phenotype_transformations.R.

#### Genomic Relatedness Matrix

We estimated the Genomic Relatedness Matrix (GRM) by calculating the realised genomic relatedness between each pair of individuals ($\pi$) using the statistical software for Genome-wide Complex Trait Analysis (GCTA(23)). Let **G** denote the GRM and let **S** denote the *n* x *m* matrix of genotypes, where *n* is the number of individuals and *m* the number of markers (i.e., SNPs), and $s_{ik}$ represents the genotypic value for an individual *i* at any SNP *k* expressed as 0,1 or 2. Then let **W** be the **S** matrix of standardised genotypic values, where any element $w_{ik}$ is:

$$w_{ik} = \frac{(s_{ik} - 2p_k)}{\sqrt{2p_k(1-p_k)}} \tag{3}$$

with $p_k$ being the MAF for the SNP *k*. Then, the GRM **G** is simply:

$$\mathbf{G} = \frac{1}{m}\mathbf{W}\mathbf{W}^{\mathbf{T}} \tag{4}$$

that is, a *n* x *n* matrix of pairwise allele frequency weighted identity by state coefficients:

$$\pi_{ij} = \frac{1}{m}\sum_{i=1}^{m}\frac{(s_{ik} - 2p_k)(s_{jk} - 2p_k)}{2p_k(1-p_k)} \tag{5}$$

where $s_{ik}$ and $s_{jk}$ are the allelic counts for any SNP *k* for the individuals *i* and *j*. The software to compute GRM can be found at https://yanglab.westlake.edu.cn/software/gcta/#MakingaGRM.

#### Threshold Genome-based Restricted Maximum Likelihood

For each trait, $h_{SNP}^2$ and $h_{\pi\geq.05}^2$ were estimated using a threshold Genome-based Restricted Maximum Likelihood approach (GREML)(24) using the GCTA software package. Consider the model:

$$\mathrm{Y} = \mathbf{X}\beta + \mathbf{W}\mathrm{u}_{\mathrm{SNP}} + \mathbf{W}_{(\pi\geq.05)}\mathrm{u}_{\pi\geq.05} + \varepsilon \tag{6}$$

where Y is the *n* x 1 vector with the proneness to feel chills score, **X** is the *n x q* covariate matrix including age, sex, genotyping array, and the first 10 Genomic PCs ($q$ = 13), β is the *q x 1* vector of fixed effects, **W** and $\mathbf{W}_{(\pi\geq.05)}$ are the matrix and spares matrix of standardised genotypes, the latter having off-diagonal elements with $\pi$ <.05 values set to 0, and $u_{SNP}$ and $u_{\pi\geq.05}$ are the vectors of random SNP and pedigree minus SNP effects. Assuming $u_{SNP} \sim N(0,\mathbf{I}\sigma_{uSNP}^2)$ and $u_{\pi\geq.05} \sim N(0, \mathbf{I}\sigma_{u\pi\geq.05}^2)$, $\sigma_Y$ (after removing β from Y) is then:

$$\sigma_Y = \mathbf{W}\mathbf{W}^{\mathbf{T}}\sigma_{uSNP}^2 + \mathbf{W}_{(\pi\geq.05)}\mathbf{W}_{(\pi\geq.05)}{}^{\mathbf{T}}\sigma_{u\pi\geq.05}^2 + \mathbf{I}\sigma_\varepsilon^2 \tag{7}$$

where **I** is the identity matrix. Since $m^*\sigma_{uSNP}^2 = \sigma_{SNP}^2$ and $m^*\sigma_{u\pi\geq.05}^2 = \sigma_{\pi\geq.05}^2$, where *m* is the number of SNPs, then $\sigma_Y$ can be rewritten as:

$$\sigma_{\mathrm{Y}} = \mathbf{G}\sigma_{\mathrm{SNP}}^2 + \mathbf{G}_{(\pi\geq.05)}\sigma_{\pi\geq.05}^2 + \mathrm{I}\sigma_\varepsilon^2 \tag{8}$$

with $\sigma_{u\pi\geq.05}^2$ capturing the difference from the pedigree-based and the SNP-based effects. By maximising the likelihood of the observed data by REML, GCTA allowed us to estimate $\sigma_{SNP}^2$, $\sigma_{u\pi\geq.05}^2$, $\sigma_{ped}^2$ (as the sum of $\sigma_{SNP}^2$ and $\sigma_{\pi\geq.05}^2$) and $\sigma_\varepsilon^2$. We note that $h_{SNP}^2$ and $h_{\pi\geq.05}^2$ are then simply calculated within GCTA as the ratio of the variance component over the total variance of Y (e.g., $h_{SNP}^2 = \sigma_{SNP}^2$ /

($\sigma_{SNP}{}^2 + \sigma_{u\pi>.05}{}^2 + \sigma_\varepsilon{}^2$)). $h_{ped}{}^2$ is provided by the GCTA software as $h_{SNP}{}^2 + h_{\pi\geq.05}{}^2$.The software to apply threshold GREML analyses can be found at https://yanglab.westlake.edu.cn/software/gcta/#GREMLinfamilydata.
Estimates obtained using a more stringent threshold of $\pi < .02$ are provided in Supplementary Table S2.

**Bivariate Threshold Genome-based Restricted Maximum Likelihood**

To estimate genetic correlations $r_g$ and $r_{\pi\geq.05}$, we used a bivariate extension of GREML. Let $Y_1$ and $Y_2$ be the two vectors with the proneness to aesthetic and music chills individuals' scores, respectively. Then:

$$Y_1 = \mathbf{X}\beta_1 + \mathbf{W}u_{SNP} + \mathbf{W}_{(\pi\geq.05)}u_{\pi\geq.05} + \varepsilon_1 \quad (9)$$

$$Y_2 = \mathbf{X}\beta_2 + \mathbf{W}u_{SNP} + \mathbf{W}_{(\pi\geq.05)}u_{\pi\geq.05} + \varepsilon_2 \quad (10)$$

Under the same set of assumptions outlined above, and under the additional assumption of no covariance between the variance components outlined below, the variance-covariance matrix **V** for $Y_1$ and $Y_1$ can be rewritten as:

$$\mathbf{V} = \mathbf{V}_{SNP} + \mathbf{V}_{\pi\geq.05} + \mathbf{V}_\varepsilon \quad (11)$$

where $\mathbf{V}_{SNP}$ is the part of the variance-covariance matrix associated with SNP effects:

$$\mathbf{V_{SNP}} = \begin{bmatrix} \sigma^2_{SNP(Y1)} & \sigma_{SNP(Y12)} \\ \sigma_{SNP(Y12)} & \sigma^2_{SNP(Y2)} \end{bmatrix} \otimes \mathbf{G}_{SNP} \quad (12)$$

$\mathbf{V}_{\pi\geq.05}$ is the part of the variance-covariance matrix associated with SNP effects in related individuals:

$$\mathbf{V}_{\pi\geq.05} = \begin{bmatrix} \sigma^2_{\pi\geq.05(Y1)} & \sigma_{\pi\geq.05(Y12)} \\ \sigma_{\pi\geq.05(Y12)} & \sigma^2_{\pi\geq.05(Y2)} \end{bmatrix} \otimes \mathbf{G}_{(\pi>.05)} \quad (13)$$

and $\mathbf{V}_\varepsilon$ is the residual part of the variance-covariance matrix:

$$\mathbf{V_e} = \begin{bmatrix} \sigma^2_{\varepsilon(Y1)} & \sigma_{\varepsilon(Y12)} \\ \sigma_{\varepsilon(Y12)} & \sigma^2_{\varepsilon(Y2)} \end{bmatrix} \otimes \mathbf{I} \quad (14)$$

with ⊗ denoting the Kronecker product. By maximising the likelihood of the observed data by (G)REML, we can estimate the variance-covariance components. $r_g$ is then calculated as:

$$r_g = \frac{\sigma_{SNP(Y12)}}{\sqrt{\sigma^2_{SNP(Y1)} * \sigma^2_{SNP(Y2)}}} \quad (15)$$

$r_{\pi\geq.05}$ is calculated similarly as $r_g$. Guidance on how to apply bivariate GREML extensions can be found at https://yanglab.westlake.edu.cn/software/gcta/#BivariateGREMLanalysis. See Supplementary Figure S1 for a graphic representation of univariate and bivariate GREML-based approaches.

**Fixed-effect inverse weighted meta-analysis of heritability estimates**

To assess the robustness of our results across the two different genotyping arrays used in Lifelines, we estimated the meta-analytic heritability obtained from fitting linear mixed models by GREML in the two subsamples independently (9244 individuals in release 1 version 2 and 6362 individuals in release 2 version 2). Within a given subpopulation and under the same environmental conditions applying to random samples of the population, let $\theta_{tp}$ be the unknown true type-specific heritability *t* (e.g., SNP) for a phenotype *p* (e.g., aesthetic chills). Then, assuming $\theta_{tp}$ to be homogeneous across the two samples of genotyped individuals, we can define $h_{tpa}^2$ as:

$$h_{\mathrm{tpa}}^2 = \theta_{tp} + \varepsilon_{tpa} \tag{16}$$

where $h_{tpa}^2$ is the estimate for $\theta_{\mathrm{tp}}$ in a sample of individuals genotyped with an array *a* (e.g., release 1, Infinium Global Screening Array®), and $\varepsilon_a \sim N(0, SE_a^2)$, with $SE_a$ being the Standard Error for the $h_{tpa}^2$. The meta-analytic estimate for $h_{\mathrm{tpa}}^2$ can then be derived as:

$$\widehat{\theta_{tp}} = \frac{\sum \mathrm{w_{tpa}} * h_{tpa}^2}{\sum w_{tpa}} \tag{17}$$

where $w_{\mathrm{tpa}} = 1/SE_{\mathrm{tpa}}^2$. Meta-analytic results can be found in Supplementary Table S4. Meta-analytic estimates were obtained using the R function rma(method = "FE") in the statistical package metafor(25). More information can be found at: https://wviechtb.github.io/metafor/reference/metafor-package.html.

**Polygenic Index-based analyses**

To estimate the association between the polygenic index (PGI) and proneness to aesthetic chills, we fit the following regression model:

$$\mathrm{y}_i = b_{PGI}\mathrm{zPGI}_i + \varepsilon_i \tag{18}$$

where *y* is the proneness to feel chills score (two-stage rank normalised and residualised data for age, sex, genotyping array, and ten genomic PCs) for the individual *i*, $b_{PGI}$ is the regression coefficient, zPGI is the standardised polygenic index for the individual *i*, and ε is the residual deviation from the predicted value for the individual *i*. Since ε cannot be assumed to be uncorrelated due to the familial relationships in the data (i.e. $\varepsilon \sim N(0, \mathbf{\Sigma})$, where $\mathbf{\Sigma} \neq \mathbf{I}*\sigma_\varepsilon^2$), we fit a model that provides robust standard errors. To do so, following reference(26), we used the sem() function in lavaan, which allows for the clustering by family via the "cluster" argument, and thus yields robust estimates of uncertainty around the point estimate.

To test whether the PGI influences differ across aesthetic and music chills, we provided a parsimonious heterogeneity PGI-based test (inspired by references(26,27)). We let η denote a phenotypic latent factor score for individual *i*, with variance defined as the covariance between the two observed traits—namely, proneness to aesthetic chills ($y_1$) and proneness to music chills ($y_2$)—such that Φ, the variance of η, is $\Phi = \mathrm{cov}(y_1, y_2)$

$$\begin{bmatrix} \mathrm{y}_{1i} \\ \mathrm{y}_{2i} \end{bmatrix} = \begin{bmatrix} 1 \\ 1 \end{bmatrix} \eta_i + \begin{bmatrix} \varepsilon_{1i} \\ \varepsilon_{2i} \end{bmatrix} \tag{19}$$

with the path coefficients from η to $y_1$ and $y_2$ fixed to unity, and regressing η on the standardised PGI, such as

$$\eta_i = b_{PGI-shared}\mathrm{zPGI}_i + \zeta_i \tag{20}$$

with $\zeta_i$ being the residual with variance Ψ. This model specification is equivalent to a bivariate common pathway model(27,28). Here, we refer to this specification as the PGI bivariate common pathway model. The model was estimated in lavaan using the sem() function, with clustering by family accounted for through the "cluster" argument. We used this model to obtain a robust $\chi^2$-distributed test statistic with 1 degree of freedom. This statistic, akin to the $Q_{trait}$ and $Q_{SNP}$, index the extent to which the effects of an exogeneous variable are not mediated by the shared variance between the two traits, with larger values indicating greater heterogeneity. Since here the exogenous variable is the PGI, we call this statistic $Q_{PGI}$. Small, non-significant $Q_{PGI}$ estimates suggest that the associations between the PGI and two traits are parsimoniously explained via shared variance. In the bivariate case, this is equivalent to testing for differences between PGI influences over $y_1$ and $y_2$. (We note that in the bivariate case, since this model has only 1 degree of freedom, there is no need to compare it with a less parsimonious model where the PGI predicts traits directly.) Since the zPGI distribution has unit variance, the overall amount of covariance explained by the PGI was calculated simply as:

$$r_{PGI-shared}{}^2 = \frac{b_{PGIs}{}^2}{b_{PGIs}{}^2 + \Psi} \tag{21}$$

A graphical representation of the common pathway PGI model can be seen in Supplementary Figure S2.

As supplementary analyses, we also provide two additional estimates for the association between the PGI and proneness to chills in a sample of unrelated to distantly related individuals ($\pi$ < .05) for transformed and raw data. The first estimate $r_{PGI(\pi < .05)}{}^2$ is obtained from fitting the model in equation 18 to the reduced sample of 10,703 individuals. The second estimate (incremental $r_{PGI(\pi < .05)}{}^2$) is obtained, in the same subset of 10,703 individuals, by fitting two models to the raw score of the proneness to chills data:

$$\mathrm{Y} = \mathbf{1_n} b + \mathbf{X} b_{cov} + \varepsilon \tag{22}$$

$$\mathrm{Y} = \mathbf{1_n} b + \mathbf{PGI} b_{PGI} + \mathbf{X} b_{cov} + \varepsilon \tag{23}$$

Where $\mathbf{1_n}$ is the n x 1 unit vector, $b$ is the intercept, **X** is the $n$ x $q$ covariate matrix including age, sex, genotyping array, and the first 10 Genomic PCs, $b_{cov}$ is the $q$ x $1$ vector of fixed effects, and the residual ε are assumed to be normally distributed such as that $\varepsilon \sim N(0, \mathbf{I}\sigma^2)$. The second model includes an $n$ x $1$ vector of PGI as well as the coefficient $b_{pgi}$. The incremental $r_{PGI(\pi < .05)}{}^2$ was obtained by subtracting the $r^2$ obtained from equation 22 with only covariates from the $r^2$ obtained from equation 23. Here, 95% Confidence Intervals (CI) were obtained by bootstrapping, resampling, with replacements, the incremental $r^2$ 1000 times.

**Expected PGI predicted accuracy**

The theoretical percentage of variance for a phenotype $y$ that can be explained by PGIs for a phenotype $x$ (derived from a genome-wide study in a completely independent sample) is:

$$r_{PGI}^2 \approx \frac{h_{SNP(x)}^2}{h_{SNP(x)}^2 + \frac{M}{N}} * h_{SNP(y)}^2 * r_G \tag{24}$$

where $h_{SNP}{}^2$ is the SNP-derived heritability for the two phenotypes, respectively, $M$ represents the number of effective SNP, $N$ is the sample size of the original genome-wide association study, and $r_g$ is the genetic correlation between $x$ and $y$(29) (see also reference(30)). Since $h_{SNP}{}^2$ for proneness to aesthetic and music chills, for openness to experience and N are known, by setting $M \sim 60{,}000$ (the approximate effective number of common SNP in European populations for common SNPs on a standard GWAS chip array(31)) we can derive the expected theoretical $r_{PGI}^2$ as a function of varying degree of $r_g$. For instance, assuming $r_g = 1$, the theoretical upper bound for $r_{PGI}^2$ equals 0.87% and 1.06%, for aesthetic and music chills, respectively. As can be seen in Supplementary Figure S3, more realistic estimates for $r_g$ would produce a smaller $r_{PGI}^2$. Equation 24 also implies that a significant $r_{PGI}^2$ is not consistent with $r_g$ between two phenotypes being equal to 0.

**Correction for assortative mating**

Estimators for $h_{SNP}^2$ are known to be upwardly biased under direct assortative mating(32). To solidify evidence of minimal upward bias on $h_{SNP}^2$ for proneness to aesthetic and music chills, we used corrected Haseman-Elston (HE) regression-based $h_{SNP}^2$ estimates ($h_{SNP\text{-}HE}^2$). We rely on the following closed-form solution for HE regression-based estimates from Border et al.(32) (as we are unaware of alternative solutions for the GREML estimator), and obtained equilibrium $h_{SNP}^2$ corrected for assortment ($h_{\infty}^2$) as:

$$h_{\infty}^2 = \frac{h_{SNP-HE}^2}{1 + r_{dAM} * h_{SNP-HE}^2} \quad (25)$$

where $h_{\infty}^2$ is the corrected heritability assuming the genetic variance has reached equilibrium in the population under study, and $r_{dAM}$ is the phenotypic partner correlation under direct assortative mating. We obtained $h_{SNP\text{-}HE}^2$ by applying the moment-based HE regressions method to aesthetic and music chills, which involves regressing the pairwise products of the (two-stage rank residualised) standardised phenotypic values ($y_i y_j$) on $\pi_{ij}$ (the latter obtained from equation 5), in other words:

$$\mathrm{y}_i \mathrm{y}_j = b_0 + b_1 \pi_{ij} + \varepsilon_{ij} \quad (26)$$

where $b_1$, the slope of such regression, is equivalent to $\sigma_{SNP}^2$. (We note, however, that HE regression is less powerful than GREML approaches(33), and that it requires removal of related individuals.) Our sample for HE regression included only 10,703 unrelated individuals (at $\pi < .05$, following(33)). The corrected $h_{\infty}^2$ were virtually indistinguishable from $h_{SNP\text{-}HE}^2$, with $h_{\infty}^2/h_{SNP\text{-}HE}^2$ ratio equal to 0.99 for aesthetic ($h_{SNP\text{-}HE}^2 = .068$ and $h_{\infty}^2 = .067$) and music chills ($h_{SNP\text{-}HE}^2 = .064$ and $h_{\infty}^2 = .063$). This indicated that assortative mating likely had little impact on $h_{SNP}^2$ estimates in this sample.

Similar to GREML analyses, HE regression was carried out through GCTA. More information can be found at https://yanglab.westlake.edu.cn/software/gcta/#Haseman-Elstonregression.

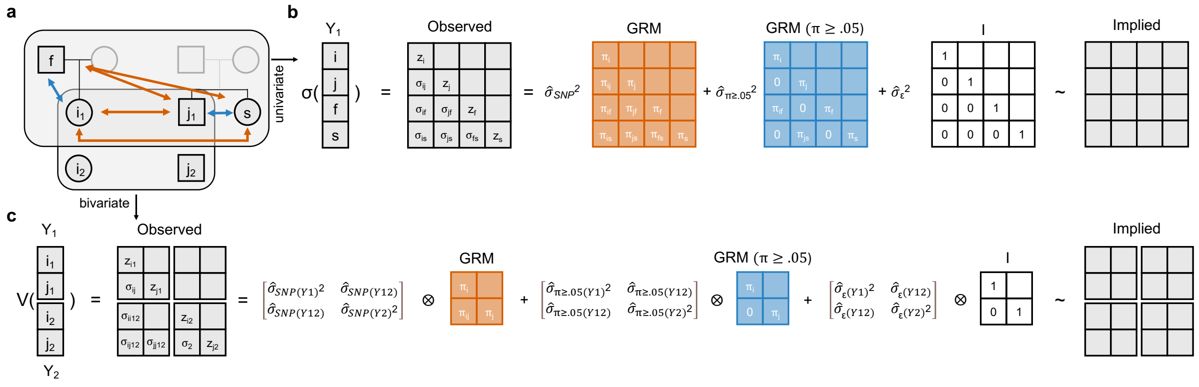


**Supplementary Figure S1.** *Conceptual representation of the GREML approach*. **a** Illustration of the relatedness structure. Coloured arrows capture the pairwise genomic relationships ($\pi$) between unrelated (warm orange, e.g., *i* and *j*) and related (azure blue, e.g., *f* and *i*) individuals. When data are available for either one ($Y_1$) or two ($Y_1$ and $Y_2$) phenotypes, information about $\pi$ can be used to estimate $h^2$ and $r_g$. **b** Univariate decomposition of the observed variance ($\sigma^2$) of $Y_1$ into different components (e.g., $\sigma_{SNP}^2$). Two Genomic Relatedness Matrices (GRM), capturing all possible pairwise $\pi$ between all and related (at $\pi \geq .05$) individuals, are used to obtain estimates for SNP contributions to phenotypic variation. Estimates are derived by maximising the likelihood of the observed data given the two GRM. Intuitively, GREML finds estimates that make the observed and the implied matrix (in grey) as similar as possible. **c** Similar to the univariate case, bivariate GREML obtains estimates for the variance-covariance components by maximising the likelihood of the observed data given the two GRM. GREML: Genome-based Restricted Maximum Likelihood Approach; $\pi$: pairwise genomic relationships; Y: phenotype; GRM: Genomic Relatedness Matrix; $\sigma_Y^2$: variance; $\sigma_{Y12}$: covariance; I: Identity matrix. Figure inspired by http://gusevlab.org/projects/hsq/.

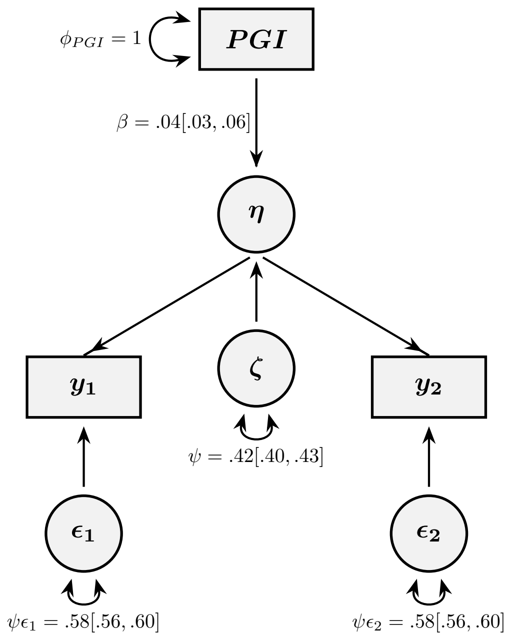


**Supplementary Figure S2.** Path diagram of the bivariate common pathway PGI model. The top rectangle represents the observed scaled PGI for openness. The bottom rectangles represent the rank-residualized scores for proneness to aesthetic chills (left) and music chills (right). The circles denote: the latent factor (η), whose variance ($B^2 * \Phi + \Psi$) corresponds to the covariance between the two traits ($cov(y_1,y_2)$); the disturbance (ζ, the residual variance Ψ of η); and the residual factors ($\varepsilon_1$ and $\varepsilon_2$). Following the literature on genomic structural equation modelling, we refer to the $\chi^2$ statistic of this model as heterogeneity Q statistic, here $Q_{PGI}$. Large $Q_{PGI}$ values indicate evidence against the null hypothesis that the PGI effect is mediated solely through the latent factor μ. Importantly, this model produces equivalent fit statistics to one in which the traits receive independent pathways with fixed coefficients, thereby providing a complementary test of whether the PGI effects on the two traits do not differ significantly.

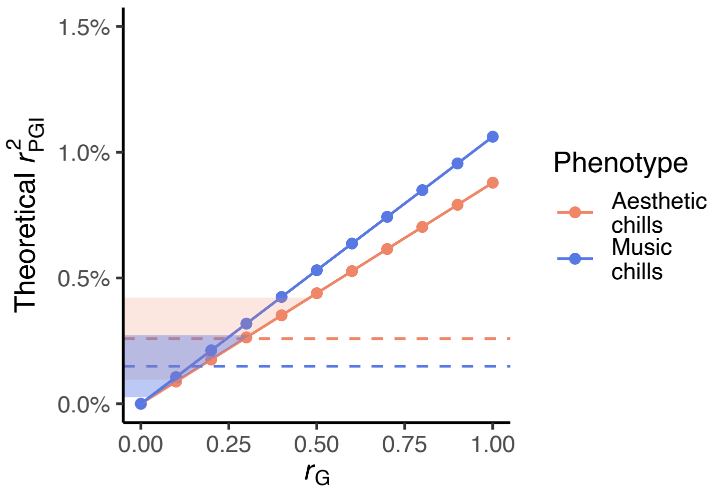


**Supplementary Figure S3.** Theoretical $r_{PGI}^2$ as a function of $r_g$. Each dot represents the theoretical expectations for the percentage of variance ($r_{PGI}^2$) in proneness to aesthetic (red) and music (blue) chills explained by the polygenic index (PGI) for openness to experience as a function of assumed genetic correlations ($r_g$) between the two traits. The dashed horizontal lines represent the observed $r_{PGI}^2$ for aesthetic (red) and music (blue) chills, with their 95% CI.

**Supplementary Table S1.** Overview of genetically informative estimates.

| Parameter | Trait | Trait status | Estimate | 95% CI |
|---|---|---|---|---|
| $h_{SNP}^2$ | Aesthetic chills | Rank transformed | .06 | [.01, .10] |
| | Music chills | Rank transformed | .07 | [.03, .11] |
| | Aesthetic chills | Raw score | .07 | [.03, .11] |
| | Music chills | Raw score | .06 | [.02, .11] |
| $h_{\pi \geq .05}^2$ | Aesthetic chills | Rank transformed | .18 | [.10, .27] |
| | Music chills | Rank transformed | .23 | [.14, .31] |
| | Aesthetic chills | Raw score | .17 | [.09, .25] |
| | Music chills | Raw score | .23 | [.14, .31] |
| $h_{ped}^2$ | Aesthetic chills | Rank transformed | .24 | [.16, .31] |
| | Music chills | Rank transformed | .29 | [.22, .36] |
| | Aesthetic chills | Raw score | .24 | [.17, .31] |
| | Music chills | Raw score | .29 | [.22 .36] |
| $r_g$ | Aesthetic-music chills | Rank transformed | .58 | [.20, 95] |
| | Aesthetic-music chills | Raw score | .58 | [.25, .92] |
| $r_{\pi \geq .05}^2$ | Aesthetic-music chills | Rank transformed | .63 | [.40, .86] |
| | Aesthetic-music chills | Raw score | .65 | [.42, .88] |
| $r_{PGI\pi < .05}^2$ | Aesthetic chills | Rank transformed | 0.3% | [0.1%, 0.5%] |
| | Music chills | Rank transformed | 0.2% | [0.0%, 0.4%] |
| $r_{PGI\pi < .05}^2$ | Aesthetic chills | Raw score | 0.2% | [0.1%, 0.4%] |
| | Music chills | Raw score | 0.1% | [0.05%, 0.3%] |

*Note.* Genetically informative results were obtained from a fully adjusted two-stage rank normalisation procedure (rank transformed) of the phenotype data(22) and raw scores (the latter including sex, age, batch, and the first ten PCs as covariates in the model). $h_{SNP}^2$: GREML-SNP-based heritability; $h_{\pi \geq .05}^2$: GREML-based excess heritability in related individuals; $h_{ped}^2$: GREML pedigree-based heritability; $r_g$: GREML-SNP-based additive genetic correlation; $r_{\pi \geq .05}^2$: GREML-SNP-based correlation between genetic components of variance in related individuals. The GREML-based estimates 95% Confidence Intervals (CI) are derived from the Standard Errors (i.e. 95% CI = estimate ± 1.96*SE). For the $r_{PGI}^2$ obtained from raw scores, $r_{PGI}^2$ 95% Confidence Intervals (CI) were obtained by bootstrapping, resampling, with replacements, the $r^2$ 1000 times.

**Supplementary Table S2.** Results of sensitivity analyses across different levels of *π.*

| Parameter | Trait | π | Estimate | 95% CI |
|---|---|---|---|---|
| $h_{SNP}^2$ | Aesthetic chills | .05 | .06 | [.01, .10] |
| | Music chills | .05 | .07 | [.03, .11] |
| | Aesthetic chills | .02 | .05 | [.01, .10] |
| | Music chills | .02 | .06 | [.02, .10] |
| $h_{\geq\pi}^2$ | Aesthetic chills | .05 | .18 | [.10, .27] |
| | Music chills | .05 | .23 | [.14, .31] |
| | Aesthetic chills | .02 | .18 | [.10, .27] |
| | Music chills | .02 | .23 | [.15, .32] |
| $h_{ped}^2$ | Aesthetic chills | .05 | .24 | [.16, .31] |
| | Music chills | .05 | .29 | [.22, .36] |
| | Aesthetic chills | .02 | .24 | [.16, .31] |
| | Music chills | .02 | .30 | [.23, .37] |
| $r_g$ | Aesthetic-music chills | .05 | .58 | [.20, 95] |
| | Aesthetic-music chills | .02 | .54 | [.15, .94] |
| $r_{\geq\pi}^2$ | Aesthetic-music chills | .05 | .63 | [.40, .86] |
| | Aesthetic-music chills | .02 | .66 | [.44, .88] |

*Note.* $h_{\geq\pi}^2$: GREML-based excess heritability in related (above or at π) individuals; $r_{\geq\pi}^2$: GREML-SNP-based correlation between genetic components of variance in related (above or at π) individuals. Other abbreviations are as in Supplementary Table S1.

**Supplementary Table S3.** Number of singletons given different genetic relatedness $\pi$

| $\pi$ | Potential relatedness | N |
|---|---|---|
| >.90 | 0th degree | 9 |
| .35 to .65 | Likely 1st degree | 2192 |
| .20 to .35 | Likely 2nd degree | 567 |
| .10 to .20 | Likely 3rd degree | 843 |
| .05 to .10 | Likely 4th degree | 1301 |
| .02 to .05 | Distant relatives | 4108 |

*Note*. 0th degree relatives, likely monozygotic twins. Likely 1st degree includes both potential full siblings or parent-offspring. Total sample size N = 15,615.

**Supplementary Table S4.** Meta-analytic heritability estimates.

| Parameter | Trait | Genotyping array | Estimate | 95% CI |
|---|---|---|---|---|
| $h_{SNP}^2$ | Aesthetic chills | Infinium Global Screening Array® | .14 | [.07, .21] |
| | Aesthetic chills | FinnGen Thermo Fisher Axiom® | .05 | [-.05, .16] |
| | Music chills | Infinium Global Screening Array® | .07 | [.00, .14] |
| | Music chills | FinnGen Thermo Fisher Axiom® | .08 | [-.02, .18] |
| $\theta_{SNP}$ | Aesthetic chills | - | .11 | [.06, .17] |
| | Music chills | - | .07 | [.02, .13] |
| $h_{\pi\geq.05}^2$ | Aesthetic chills | Infinium Global Screening Array® | .09 | [-.03, .22] |
| | Aesthetic chills | FinnGen Thermo Fisher Axiom® | .20 | [.02, .39] |
| | Music chills | Infinium Global Screening Array® | .24 | [.13, .36] |
| | Music chills | FinnGen Thermo Fisher Axiom® | .14 | [-.03, .32] |
| $\theta_{\pi\geq.05}$ | Aesthetic chills | - | .13 | [.03, .23] |
| | Music chills | - | .21 | [.12, .31] |
| $h_{ped}^2$ | Aesthetic chills | Infinium Global Screening Array® | .24 | [.14, .34] |
| | Aesthetic chills | FinnGen Thermo Fisher Axiom® | .26 | [.11, .41] |
| | Music chills | Infinium Global Screening Array® | .32 | [.22, .41] |
| | Music chills | FinnGen Thermo Fisher Axiom® | .22 | [.07, .37] |
| $\theta_{ped}$ | Aesthetic chills | - | .24 | [.16, .33] |
| | Music chills | - | .29 | [.21, .37] |

*Note.* $h_{SNP}^2$: GREML-SNP-based heritability; $\theta_{SNP}$: meta-analytic estimate for $h_{SNP}^2$; $h_{\pi\geq.05}^2$: GREML-based excess heritability in related individuals. $\theta_{\pi\geq.05}$: meta-analytic estimate for $h_{\pi\geq.05}^2$. $h_{ped}^2$: GREML pedigree-based heritability. $\theta_{ped}$: meta-analytic estimate for $h_{\pi\geq.52}^2$. For GREML-based estimators, the 95% Confidence Intervals (CI) are derived from the Standard Errors (i.e. 95% CI = estimate ± 1.96*SE), for meta-analytic estimates 95% CI are derived directly from the rma(method = "FE") output.

**List of Legends**

**Supplementary Figure S1.** *Conceptual representation of the GREML approach*. **a** Illustration of the relatedness structure. Coloured arrows capture the pairwise genomic relationships ($\pi$) between unrelated (warm orange, e.g., *i* and *j*) and related (azure blue, e.g., *f* and *i*) individuals. When data are available for either one ($Y_1$) or two ($Y_1$ and $Y_2$) phenotypes, information about $\pi$ can be used to estimate $h^2$ and $r_g$. **b** Univariate decomposition of the observed variance ($\sigma^2$) of $Y_1$ into different components (e.g., $\sigma_{SNP}^2$). Two Genomic Relatedness Matrices (GRM), capturing all possible pairwise $\pi$ between all and related (at $\pi \geq .05$) individuals, are used to obtain estimates for SNP contributions to phenotypic variation. Estimates are derived by maximising the likelihood of the observed data given the two GRM. Intuitively, GREML finds estimates that make the observed and the implied matrix (in grey) as similar as possible. **c** Similar to the univariate case, bivariate GREML obtains estimates for the variance-covariance components by maximising the likelihood of the observed data given the two GRM. GREML: Genome-based Restricted Maximum Likelihood Approach; $\pi$: pairwise genomic relationships; Y: phenotype; GRM: Genomic Relatedness Matrix; $\sigma_Y^2$: variance; $\sigma_{Y12}$: covariance; I: Identity matrix. Figure inspired by http://gusevlab.org/projects/hsq/.

**Supplementary Figure S2.** Path diagram of the bivariate common pathway PGI model. The top rectangle represents the observed scaled PGI for openness. The bottom rectangles represent the rank-residualized scores for proneness to aesthetic chills (left) and music chills (right). The circles denote: the latent factor (η), whose variance ($B^2 * \Phi + \Psi$) corresponds to the covariance between the two traits ($cov(y_1,y_2)$); the disturbance (ζ, the residual variance Ψ of η); and the residual factors ($\varepsilon_1$ and $\varepsilon_2$). Following the literature on genomic structural equation modelling, we refer to the $\chi^2$ statistic of this model as heterogeneity Q statistic, here $Q_{PGI}$. Large $Q_{PGI}$ values indicate evidence against the null hypothesis that the PGI effect is mediated solely through the latent factor µ. Importantly, this model produces equivalent fit statistics to one in which the traits receive independent pathways with fixed coefficients, thereby providing a complementary test of whether the PGI effects on the two traits do not differ significantly.

**Supplementary Figure S3.** Theoretical $r_{PGI}^2$ as a function of $r_g$. Each dot represents the theoretical expectations for the percentage of variance ($r_{PGI}^2$) in proneness to aesthetic (red) and music (blue) chills explained by the polygenic index (PGI) for openness to experience as a function of assumed genetic correlations ($r_g$) between the two traits. The dashed horizontal lines represent the observed $r_{PGI}^2$ for aesthetic (red) and music (blue) chills, with their 95% CI.

**Supplementary Table S1.** Overview of genetically informative estimates. *Note.* Genetically informative results were obtained from a fully adjusted two-stage rank normalisation procedure (rank transformed) of the phenotype data(22) and raw scores (the latter including sex, age, batch, and the first ten PCs as covariates in the model). $h_{SNP}^2$: GREML-SNP-based heritability; $h_{\pi\geq.05}^2$: GREML-based excess heritability in related individuals; $h_{ped}^2$: GREML pedigree-based heritability; $r_g$: GREML-SNP-based additive genetic correlation; $r_{\pi\geq.05}^2$: GREML-SNP-based correlation between genetic components of variance in related individuals. The GREML-based estimates 95% Confidence Intervals (CI) are derived from the Standard Errors (i.e. 95% CI = estimate ± 1.96*SE). For the $r_{PGI}^2$ obtained from raw scores, $r_{PGI}^2$ 95% Confidence Intervals (CI) were obtained by bootstrapping, resampling, with replacements, the $r^2$ 1000 times.

**Supplementary Table S2.** *Note.* $h_{\geq\pi}^2$: GREML-based excess heritability in related (above or at π) individuals; $r_{\geq\pi}^2$: GREML-SNP-based correlation between genetic components of variance in related (above or at π) individuals. Other abbreviations are as in Supplementary Table S1.

**Supplementary Table S3.** Number of singletons given different genetic relatedness *π. Note*. 0th degree relatives, likely monozygotic twins. Likely 1st degree includes both potential full siblings or parent-offspring. Total sample size N = 15,615.

**Supplementary Table S4.** Meta-analytic heritability estimates. *Note.* $h_{SNP}^2$: GREML-SNP-based heritability; $\theta_{SNP}$: meta-analytic estimate for $h_{SNP}^2$; $h_{\pi\geq.05}^2$: GREML-based excess heritability in related individuals. $\theta_{\pi\geq.05}$: meta-analytic estimate for $h_{\pi\geq.05}^2$. $h_{ped}^2$: GREML pedigree-based heritability. $\theta_{ped}$: meta-analytic estimate for $h_{\pi\geq.52}^2$. For GREML-based estimators, the 95% Confidence Intervals (CI) are derived from the Standard Errors (i.e. 95% CI = estimate ± 1.96*SE), for meta-analytic estimates 95% CI are derived directly from the rma(method = “FE”) output.
